# Parabrachial Calcitonin Gene-Related Peptide Neurons Sex-Specifically Modulate Anxiety in Alcohol Withdrawal

**DOI:** 10.1101/2025.11.24.690162

**Authors:** C.E. Van Doorn, J.B. Tyree, A.A. Jaramillo

## Abstract

Parabrachial nucleus (PBN) neurons expressing Calcitonin Gene-Related Peptide (CGRP) modulate fear– and anxiety-like behavior. It is unknown if PBN(CGRP) neurons play a role in anxiety during withdrawal from alcohol or after repeated stress. First, to investigate the anxiogenic role of activating PBN(CGRP) neurons in naïve conditions, *Calca*^CRE^ female and male mice expressing CRE-dependent hM3D(Gq) DREADDs in the PBN were tested on the elevated plus maze (EPM). PBN(CGRP) neurons drive phasic activity in the bed nucleus of the stria terminalis (BNST) that synchronizes to anxiety-like behavior. Therefore, a transsynaptic anterograde AAV-based strategy was used in C57BL/6J female and male mice to activate BNST neurons innervated by the PBN projections (BNST^PBN^) during EPM. Additionally, PBN(CGRP) neurons and CGRP-innervated BNST cells were measured in experimentally-naïve CGRP-DTR^GFP^ female and male mice, to investigate baseline sex-differences. Next, to investigate the impact of PBN(CGRP) inhibition on anxiety-like behavior following chronic intermittent ethanol vapor exposure (CIE), *Calca*^CRE^ female and male mice expressing CRE-dependent hM4D(Gi) DREADDs in the PBN were tested in EPM during acute withdrawal. Additionally, mice were exposed to repeated forced swim stress (FSS) paired with PBN(CGRP) inhibition followed by testing in the novelty suppressed feeding task (NSFT), to investigate the role of PBN(CGRP) on anxiety-like behavior after stress in prolonged withdrawal. Activating PBN(CGRP) or BNST^PBN^ neurons did not change behavior in EPM in either sex. Total PBN(CGRP) and CGRP-innervated BNST cells did not differ between females and males. In acute withdrawal, inhibiting PBN(CGRP) neurons decreased time in the open arm and increased time in the closed arm selectively in females. In prolonged withdrawal, inhibiting PBN(CGRP) neurons during FSS did not affect immobility in either sex, but did subsequently decrease approach frequency and average speed selectively in males. Altogether, the data report the anxiogenic effect of manipulating PBN(CGRP) neurons is stress and sex-specific during alcohol withdrawal.

## 2. Introduction

Despite the prevalence of Alcohol Use Disorder (AUD), there are limited therapeutic options. A characteristic of AUD is the presence of negative affect during withdrawal, which increases the likelihood of relapse (1, 2). Moreover, stressful events occurring during withdrawal can also decrease the likelihood of sobriety (1, 2). Neuropeptide systems are drug target candidates for treating AUD due to their potential role in negative affect– and stress-related relapse. The neuropeptide Calcitonin Gene-Related Peptide (CGRP) modulates anxiety-like behavior, pain, and fear (3, 4). Monoclonal antibodies targeting the CGRP system are shown to be safe and an effective treatment for migraines in humans (e.g. epitinezumab, erenumab; (5)) and are currently being considered for the treatment of hangover migraines (6, 7). Given CGRP monoclonal antibodies are well-tolerated and bioavailable in humans, inhibiting CGRP is a promising neuropeptide target for an AUD pharmacotherapy.

CGRP has established roles in anxiety and stress contexts (3, 8–11). Blocking CGRP signaling peripherally, intracerebroventricularly, or site-specifically in the bed nucleus of the stria terminalis (BNST), a critical node for anxiety (3, 12), alters stress responses and decreases anxiety-like behavior (8–11, 13–16). Neurons in the parabrachial nucleus (PBN) are the predominate source of CGRP in the BNST (3, 4, 16–18). Similar to CGRP modulation, activating PBN neurons that express CGRP (PBN[CGRP]) increased threat responses such as fear– and anxiety-like behavior in both sexes (3, 4, 19–22). Additionally, we have demonstrated that PBN(CGRP) neurons drive BNST activity that synchronized to anxiety-like behavior (21). Interestingly, circuit-specific modulation reveals a sex-specific role in anxiety, as activating BNST neurons that are innervated by PBN projections (BNST^PBN^) selectively increased anxiety-like behavior in females (21). In males, the role of the PBN(CGRP) ➜ BNST circuit in anxiety may occur after stress exposure, as neurotransmission driven by PBN(CGRP) projections in the BNST is dysregulated weeks after a stress-induced anxiety-like state (23). Thus, in addition to driving anxiety-like behavior, exposure to stress and negative affective states may have long-term effects on the PBN(CGRP) ➜ BNST circuit. Given the potential role of PBN(CGRP) neurons in negative affect, modulating PBN(CGRP) signaling may be a viable target for the treatment of AUD.

Although limited, the current data suggest a relationship between CGRP and alcohol. A clinical study in patients with psoriasis demonstrates a polymorphism in the *Calca* gene, which encodes for CGRP, was associated with non-drinking patients compared to patients with a history of drinking (24). Preclinical studies also suggest a genetic predisposition to alcohol that is associated with the CGRP system, as in alcohol-naïve conditions alcohol-preferring rats demonstrated differential CGRP receptor and peptide expression in select brain regions (25, 26). Additionally, CGRP terminal-expression in the BNST changed with voluntary alcohol exposure in alcohol-preferring rats in a sub-region specific manner (27). The effect of alcohol on the CGRP system extends to withdrawal, as weeks of withdrawal from voluntary drinking increased CGRP immunoreactivity in select brain regions (27). Like stress, alcohol induces long-term changes in PBN(CGRP) circuit activity, as activating the BNST in adult female mice with a history of adolescent alcohol vapor exposure increased activity in PBN(CGRP) neurons (28). Given alcohol induces changes in PBN(CGRP) activity, and its activation is anxiogenic, CGRP may have a role in anxiety during alcohol withdrawal.

First, to determine the role of the PBN(CGRP) and BNST^PBN^ neurons on anxiety-like behavior in alcohol-naïve conditions, we focus on behavior during the elevated plus maze (EPM). To determine if there are sex-differences in PBN(CGRP) neurons and CGRP-innervated BNST neurons, we quantified CGRP neuron expression in the PBN and fiber expression in the BNST. Next, we utilized chronic intermittent ethanol vapor exposure (CIE) and EPM to determine the impact of PBN(CGRP) inhibition on anxiety-like behavior in acute withdrawal. To evaluate the anxiolytic potential of repeated PBN(CGRP) inhibition during stress in prolonged abstinence from CIE, we utilized repeated forced swim stress (FSS) and novelty suppressed feeding task (NSFT), two paradigms in which behavior has been shown to be modulated by CGRP or PBN(CGRP) neurons (8, 21, 23). We hypothesize that activating PBN(CGRP) and BNST^PBN^ neurons drives anxiety-like responses in alcohol-naïve mice, and inhibiting PBN(CGRP) in early alcohol withdrawal decreases anxiety-like behaviors. Additionally, we hypothesize that inhibiting PBN(CGRP) activity during stress exposure in prolonged abstinence decreases anxiety-like behaviors. We expect to find sex-differences in PBN(CGRP)-modulated behavior and CGRP expression. Overall, we expect our findings to elucidate the role of PBN(CGRP) neurons in anxiety during early and prolonged alcohol withdrawal.

## 3. Materials and Methods

### 3.1. Animals

Female and male *Calca*^CRE^ mice (B6.Cg-Calcatm1.1[cre/EGFP]Rpa/J) were used for Experiments 1, 4 and 5. *Calca*^CRE^ mice are a heterozygous genetic knock-in mouse model that express CRE-recombinase at the *Calca* locus with a C57BL/6J background (29). For experiment 2, female and male C57BL/6J mice were used. For experiment 3, female and male CGRP-DTR^GFP^ mice (Calcitonin gene-related peptide [cre/EGFP] (CGRPalpha DTR) were used. CGRP-DTR^GFP^ knock-in mice express the axonal tracer (farnesylated enhanced green fluorescent protein; GFP) and LoxP-stopped cell ablation construct (human diphtheria toxin receptor; hDTR) to the *Calca* locus (30). Both transgenic lines were bred in-house and genotyped with Transnetyx (Transetyx, Inc, Cordova, TN, United States).

All mice were 10-12 weeks old at the start of experiments. Mice were single-(Experiment 1) or group-housed (Experiments 2-5) with water and food available ad lib in the home cage. The colony room was maintained on a 12-h light/dark cycle (lights on at 06:00) under controlled temperature (20–25 °C) and humidity (30–50%) levels. All experiments were conducted during the light phase, with the exception of CIE, which was conducted in the dark phase. Animals were under continuous care and monitoring by veterinary staff from the Vanderbilt Division of Animal Care or University of Kentucky Division of Lab Animal Resources. All procedures were carried out in accordance with the NIH Guide to Care and Use of Laboratory Animals and institutional guidelines and approved by the Institutional Animal Care and Use Committee at Vanderbilt University and the University of Kentucky.

### 3.2. Stereotaxic viral surgeries

For Experiments 1-2, 4 and 5, adult mice were anesthetized with isoflurane (Covetrus, Portland, Maine, United States; initial dose = 3%; maintenance dose = 1.5%) for surgery performed using a stereotax (Leica Biosystems, Nussloch, Germany or RWD Life Science, Shenzhen, Guangdong, PR, China). Coordinates from bregma were used to target the PBN (AP = −5.34, ML±1.31, DV = −3.37, 15.03° angle). Additionally, for Experiment 2 the BNST was targeted (AP = 0.14, ML ± 0.88, DV = −4.18, 15.03° angle). Care was taken to prevent drying of the eye by applying artificial tears ocular lubricant (Akorn, Lake Forest, IL, United States) and reapplying as needed throughout the surgery. Viral vectors were infused bilaterally at 300 nL (40 nL/min) with the needle remaining in place for an additional 5-min before withdrawal. Experiment 1 mice received a CRE-dependent virus expressing excitatory DREADDs (AAV5-hSyn-DIO-hM3D[Gq]-mCherry, hM3D[Gq]; Addgene, Watertown, MA, USA) in the PBN. Experiment 2 mice received a virus expressing CRE (AAV1-hSyn-CRE; Addgene, Watertown, MA, USA) in the PBN and a CRE-dependent virus expressing excitatory DREADDs (AAV5-hSyn-DIO-hM3D[Gq]-mCherry, hM3D[Gq]; Addgene, Watertown, MA, USA) in the BNST. For Experiments 4 and 5, mice received a CRE-dependent virus expressing inhibitory DREADDs (AAV5-hSyn-DIO-hM4D[Gi]-mCherry, hM4D[Gi]; Addgene, Watertown, MA, USA). Mice were given 0.9% saline for fluid maintenance postoperatively and body weights were tracked daily to ensure body weight was maintained. Mice were treated with Alloxate (Pivetal, Liberty, MO, United States; 2.5 mg/kg, S.C.) or Meloxicam (Covetrus, Portland, Maine, United States; 2.5mg/kg S.C.) for 72-hr after surgery and allowed to recover for a 1 week. To allow for sufficient DREADD expression, mice received chemogenetic manipulations after >4 weeks. Viral deposits were confirmed post-testing utilizing autofluorescence or immunofluorescence protocols described below.

### 3.3. Immunofluorescence and Viral Vector Confirmation

For all experiments, mice were deeply anesthetized using isoflurane, transcardially perfused with ice-cold phosphate buffered saline (PBS) followed by 4% paraformaldehyde (PFA) in PBS. Brains were submerged in 4% PFA for 24-hr at 4 °C and cryoprotected in 30% sucrose in PBS for a minimum of 5 days. Coronal sections were cut on a cryostat (Leica CM3050S, Leica Biosystems, Nussloch, Germany) in Optimal Cutting Temperature solution (VWR, Radnor, PA, United States) at a thickness of 40 μm and stored in PBS at 4 °C until immunofluorescence or autofluorescence analysis. To visualize CGRP expression for Experiment 3, sections were incubated in goat anti-CGRP primary antibody (1:400; AB36001, Abcam, Waltham, MA, USA) for 48-hr at room temperature, washed in PBS, and incubated in Cy3 donkey anti-goat secondary antibody (1:500; Cy3 AffiniPure Donkey Anti-Goat, 705-165-003, Jackson, West Grove, PA, USA) in 0.1% Triton X-100 in PBS for 24-hr at 4 °C and incubated in DAPI (1:10,000; 17509, AAT Bioquest, Sunnyvale, CA, USA) in de-ionized water. Sections were washed with PBS and mounted on Fisher Plus slides (Fisher Scientific, Waltham, MA, USA) and cover slipped with Poly AquaMount (Polysciences, Warrington, PA, USA) when dry. Images were taken on Nikon Multi Excitation TIRF microscope (Nikon Metrology Inc, Tokyo, Japan) at 10x and 20x magnification. ImageJ software (NIH) was used to assess the total number of co-localized CGRP+ and DAPI+ cells in the PBN and CGRP+ innervated DAPI+ cells in the BNST. For viral placement confirmation, in Experiment 1, hM3D(Gq) expression was confirmed by imaging mCherry autofluorescence in the PBN with a Keyence BZ-X710 microscope (Keyence, Itasca, IL, United States) at 20x magnification. For Experiment 2, hM3D(Gq) expression was confirmed by imaging mCherry autofluorescence in the BNST with Nikon Multi Excitation TIRF microscope (Nikon Metrology Inc, Tokyo, Japan) at 10x magnification. For Experiment 4-5, hM4D(Gi) expression was confirmed by imaging mCherry autofluorescence in the PBN with ECHO Revolve microscope (ECHO, San Diego, CA, United States) at 20x magnification.

### 3.4. Chronic Intermittent Ethanol (CIE)

Mice underwent CIE described in (23, 31). Briefly, mice received a daily intraperitoneal (IP) injection of pyrazole (1 mmol/kg) combined with 1.6 g/kg ethanol and were placed in an ethanol vapor (95% ethanol at 5 L/min) enclosure (VIPER system, LJari, La Jolla, CA, United States) for 16 hours per day, then returned to standard animal housing for 8 hours. This was repeated over the course of four days, followed by a three-day stay in regular animal housing, and then another four days of the exposure process (i.e., two CIE cycles).

### 3.5. Behavioral Testing

For all testing, mice were brought into the testing room and allowed to acclimate for one hour. Mice were visualized, recorded, and tracked by an overhead camera using AnyMaze software (Stoelting Co, Wood Dale, IL, United States). Behavioral analyses were conducted by an observer blinded to treatment conditions.

#### 3.5.1. Elevated Plus Maze (EPM)

The maze is elevated 55 cm above the ground and consists of two open arms and two closed arms (30.5 × 6.5 cm; 16 cm closed arm height) with a 5 × 5 cm open center zone. Lux was measured at the center of each open arm at 85-105 lux, center at 70 lux, and each closed arm at 0-6 lux by using a LX1010B Digital Light Meter (FTVOGUE, China). Mice were placed in the center facing alternate arms and recorded for 5 min.

#### 3.5.2. Forced Swim Stress (FSS)

FSS was performed as previously described (23). Briefly, mice were placed into a transparent cylindrical container (i.e., 5000 mL glass beaker) with water at room temperature (20.6 – 21.1° C) for 6 min. Lux was measured at the center of the surface of the water at 300-350 lux using a LX1010B Digital Light Meter (FTVOGUE, China). Water was changed in between animals. The last 4 min of the test were analyzed for immobility.

#### 3.5.3. Novelty-Suppressed Feeding Test (NSFT)

The 3-day NSFT was performed as previously described (21, 23, 32). Briefly, mice were food restricted for 48 h except for 2 h food access 22-24 h before testing. The testing apparatus was a 50 x 50 cm arena with fresh home cage bedding and a single chow pellet at the center. Lux was measured at the center of the arena at 300-450 lux with a LX1010B Digital Light Meter (FTVOGUE, China). At the start of testing, mice were placed in a corner of the arena. Food approach frequency was defined as approach to the 10 cm diameter around the center-located food pellet. Mice were removed from the testing arena immediately after the first bite (i.e., consummatory food approach) and placed in their home cage with the food pellet from the test. The pellet was weighed after 10 min to confirm food consumption. Mice were returned to ad lib access to chow post testing. Latency to the first bite and food consumption were hand-scored by an observer blinded to treatment groups.

### 3.6. Experimental Procedures

#### 3.6.1. Experiment 1: Effects of PBN(CGRP) activation on EPM

*Calca*^CRE^ female (n=9) and male mice (n=7) received bilateral injections of CRE-dependent virus to express hM3D(Gq) in the PBN, as previously reported in (21). Mice with inefficient viral expression (2 females) received saline treatment and therefore were included in the data. Mice received Clozapine-N-oxide (CNO, 3 mg/kg, intraperitoneal, I.P.) or saline in their home cage 30-min prior to the start of EPM testing (**Figures 1**-**2**).

**Figure 1.**
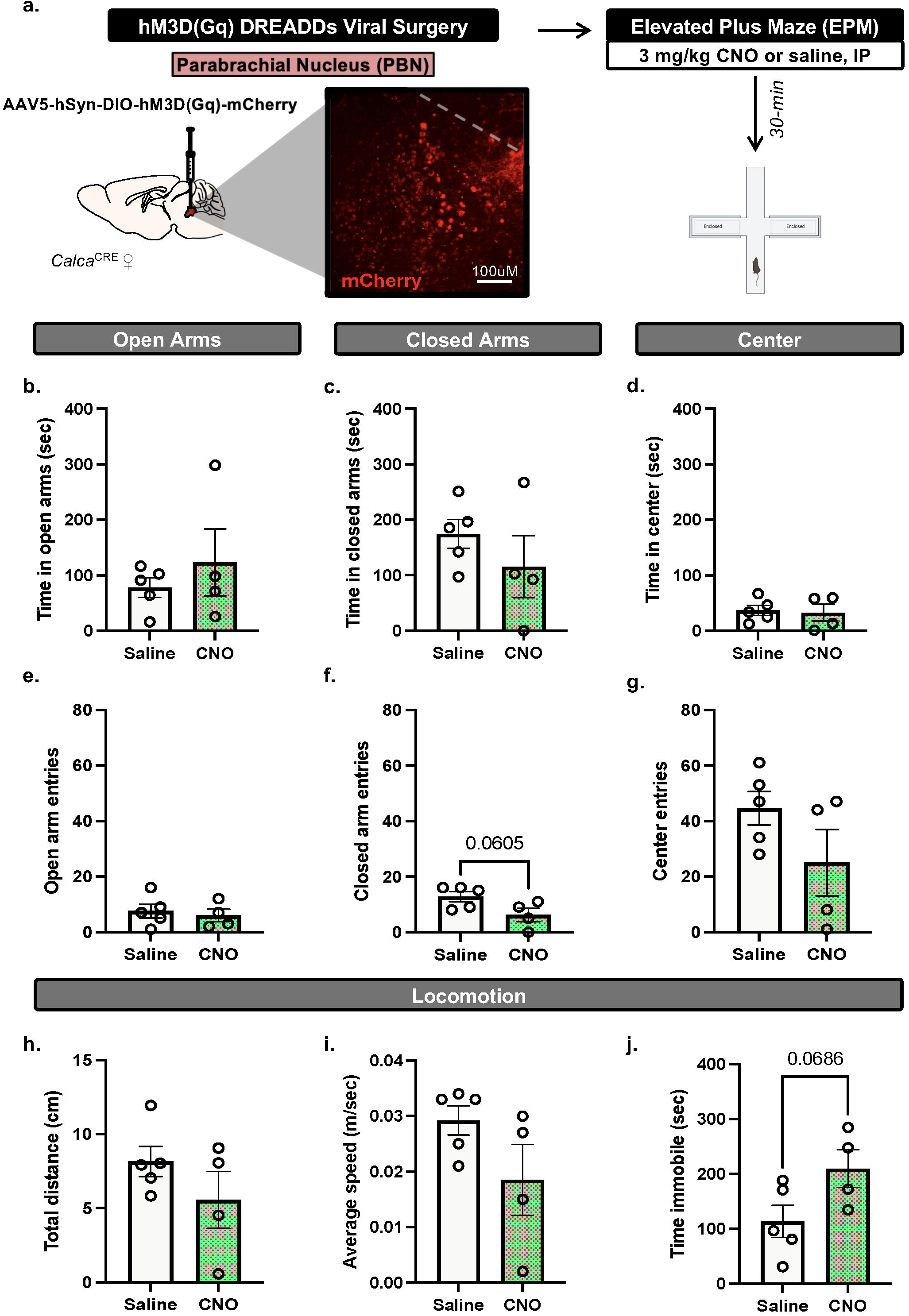
PBN(CGRP) activation does not change EPM behaviors in females. **a)** *Calca*^Cre^ female mice received bilateral infusions to express CRE-dependent hM3D(Gq) DREADDs in the PBN and were tested with EPM post-30 min pretreatment of CNO (3mg/kg, IP). Inset demonstrates representative image of hM3D(Gq) (mCherry) autofluorescence in the PBN. **b**) CNO did not change time in open arms, **c**) time in closed arms, **d**) time in center, **e**) open arm entries, **f**) closed arm entries, **g**) center entries, **h)** distance travelled, i) average speed or **j**) time immobile in EPM compared to saline. Values on graphs represent mean ± S.E.M., unpaired *t*-test (*p≦0.05).

**Figure 2.**
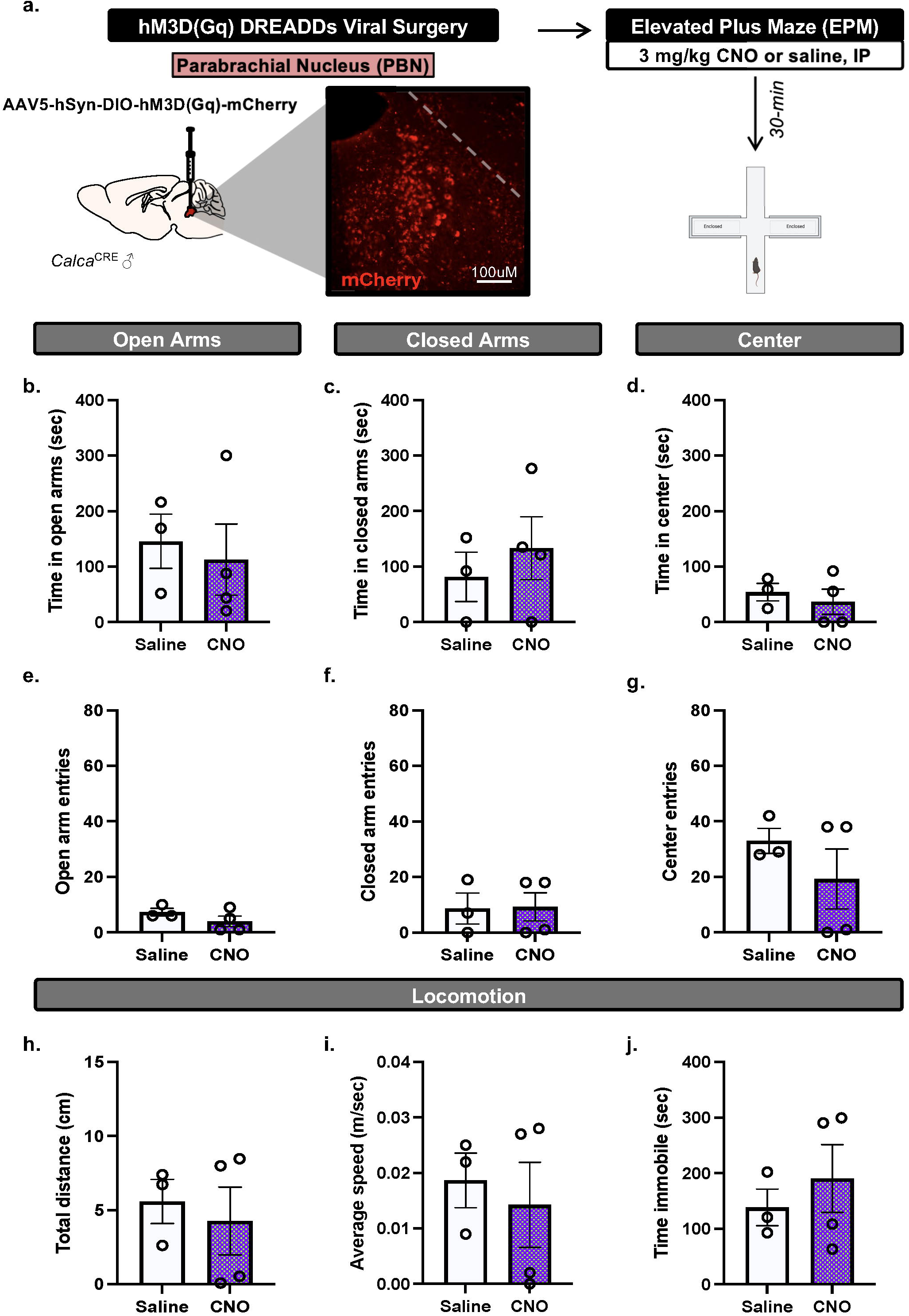
PBN(CGRP) activation does not change EPM behaviors in males. **a)** *Calca*^Cre^ male mice received bilateral viral infusions to express CRE-dependent hM3D(Gq) DREADDs in the PBN and were tested with EPM post-30 min pretreatment of CNO (3mg/kg, IP). Inset demonstrates representative image of hM3D(Gq) (mCherry) autofluorescence in the PBN. **b)** CNO did not change time in open arms, **c**) time in closed arms, **d**) time in center, **e**) open arm entries, **f**) closed arm entries, **g**) center entries, **h**) distance travelled, **i**) average speed or **j**) time immobile in EPM compared to saline. Values on graphs represent mean ± S.E.M., unpaired *t*-test (*p≦0.05).

#### 3.6.2. Experiment 2: Effects of BNST^PBN^ activation on EPM

C57BL/6J female (n=8) and male mice (n=7) received bilateral injections of the transsynaptic anterograde CRE-expressing virus in the PBN and CRE-dependent virus in the BNST. Thereby, hM3D(Gq) would be expressed in BNST neurons that receive projections from the PBN, as previously reported in (21). Mice with inefficient viral expression (3 females) were not included in the data. Mice received CNO (3 mg/kg, intraperitoneal, I.P.) or saline in their home cage 30-min prior to the start of EPM testing (**Figures 3-4**). 3 weeks later mice were re-tested with EPM albeit with the opposite treatment (e.g., mice that received CNO for the first EPM test now received saline).

**Figure 3.**
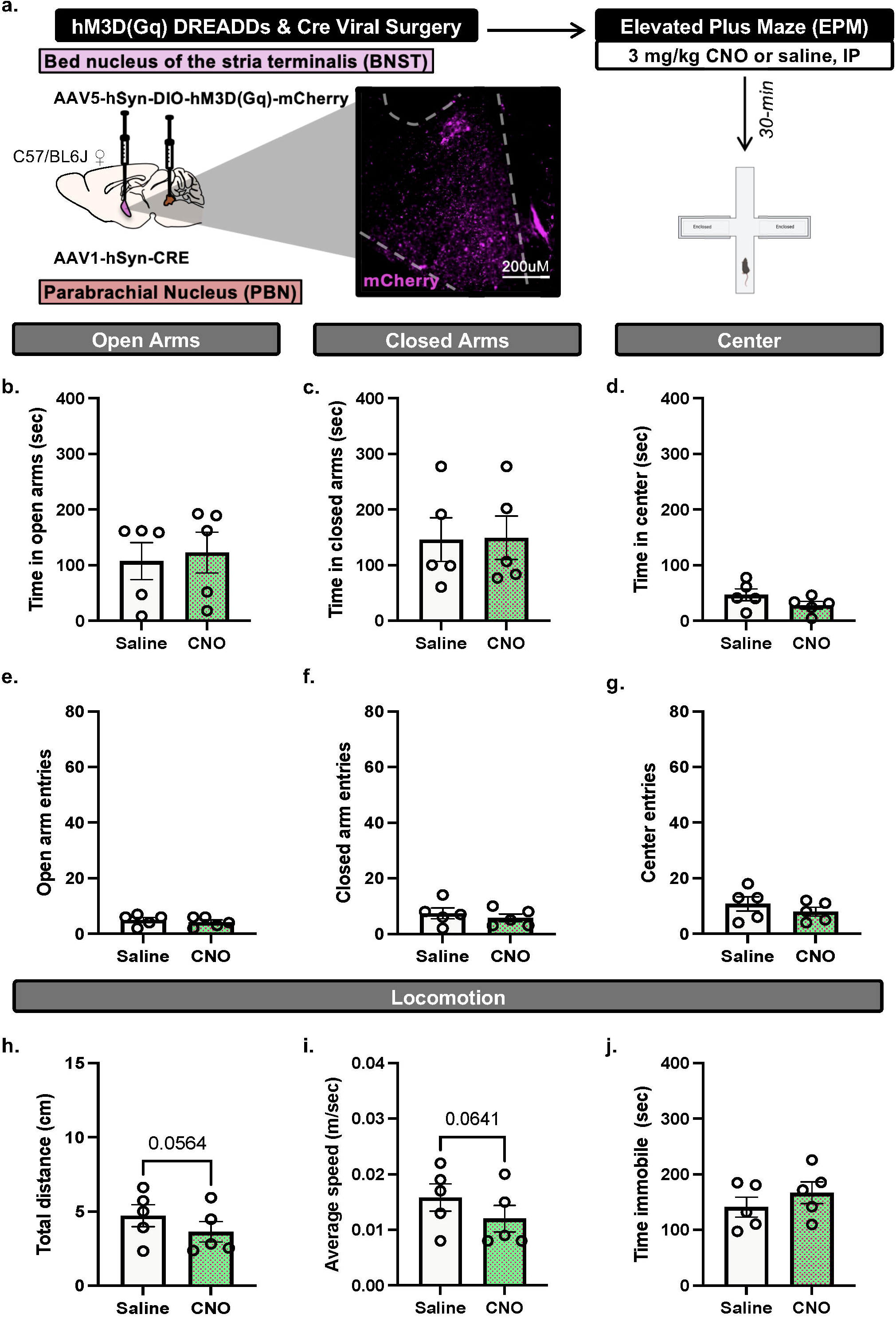
BNST^PBN^ activation does not change EPM behaviors in females. **a)** *Calca*^Cre^ female mice received bilateral infusions of AAV1-CRE anterograde transsynaptic virus in the PBN and CRE-dependent hM3D(Gq) DREADDs virus in the BNST, to express hM3D(Gq) DREADDs in BNST neurons that are innervated by PBN projections. Mice were tested with EPM post-30 min pretreatment of CNO (3mg/kg, IP). Inset demonstrates representative image of hM3D(Gq) (mCherry) autofluorescence in the BNST. **b**) CNO did not change time in open arms, **c**) time in closed arms, **d**) time in center, **e**) open arm entries, **f**) closed arm entries, **g**) center entries, **h**) distance travelled, **i**) average speed or **j**) time immobile in EPM compared to saline. Values on graphs represent mean ± S.E.M., paired *t*-test (*p≦0.05).

**Figure 4.**
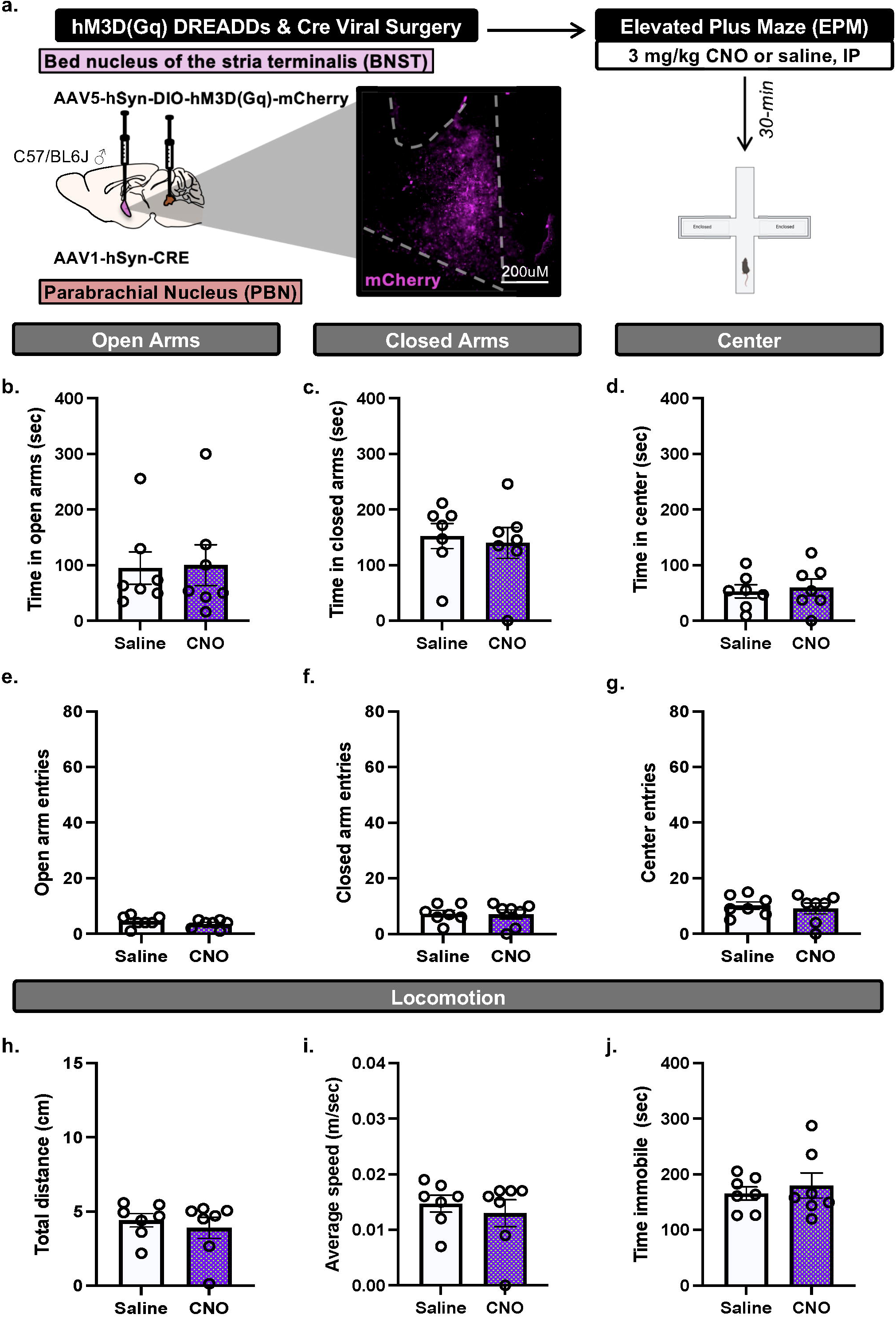
BNST^PBN^ activation does not change EPM behaviors in males. **a)** *Calca*^Cre^ male mice received bilateral infusions of AAV1-CRE anterograde transsynaptic virus in the PBN and CRE-dependent hM3D(Gq) DREADDs virus in the BNST, to express hM3D(Gq) DREADDs in BNST neurons that are innervated by PBN projections. Mice were tested with EPM post-30 min pretreatment of CNO (3mg/kg, IP). Inset demonstrates representative image of hM3D(Gq) (mCherry) autofluorescence in the BNST. **b**) CNO did not change time in open arms, **c**) time in closed arms, **d**) time in center, **e**) open arm entries, **f**) closed arm entries, **g**) center entries, **h**) distance travelled, **i**) average speed or **j**) time immobile in EPM compared to saline. Values on graphs represent mean ± S.E.M., paired t-test (*p≦0.05).

#### 3.6.3. Experiment 3: CGRP Expression in PBN neurons and fibers in the BNST

CGRP-DTR^GFP^ female (n=6) and male (n=6) mice were experimentally naïve (i.e. no behavioral assay exposure). PBN-containing slices were analyzed for co-localization of CGRP (GFP) and DAPI and reported as total co-localized (**Figure 5a-b**). BNST-containing slices were analyzed for co-localization of DAPI innervated by CGRP (GFP) expression and reported as total co-localized (**Figure 5c-d**).

**Figure 5.**
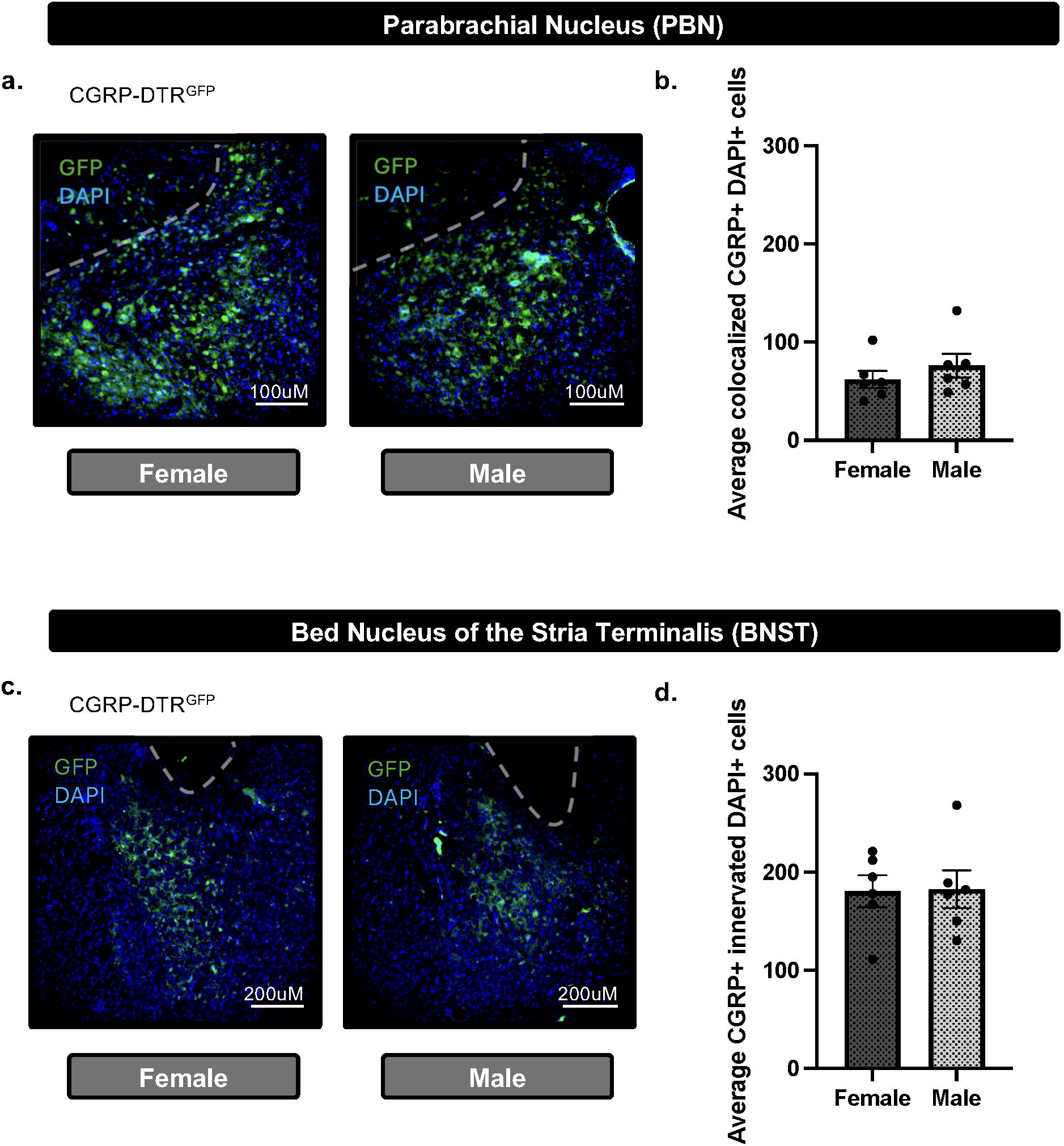
CGRP+ cells in the PBN and CGRP+ innervated cells in the BNST do not differ between females and males. **a)** Representative image demonstrating CGRP (GFP) and DAPI merged immunofluorescence in the PBN of CGRP-DTR^GFP^ females and males. **b**) Quantification of CGRP (GFP) demonstrate no difference in total CGRP+DAPI colocalization in the PBN between males and females. **c**) Representative image demonstrating CGRP (GFP) and DAPI merged immunofluorescence in the BNST of CGRP-DTR^GFP^ females and males. **d**) Quantification of CGRP (GFP) demonstrate no difference in total CGRP-innervated DAPI+ colocalization in the PBN between males and females. Values on graphs represent mean ± S.E.M., unpaired *t*-test (*p≦0.05).

#### 3.6.4. Experiment 4: Effects of PBN(CGRP) inactivation on EPM during acute CIE withdrawal

*Calca*^CRE^ female (n=7) and male (n=10) mice received CRE-dependent viral injection to express hM4D(Gi) in the PBN. Mice with inefficient viral expression (2 female, 6 males) were included in the data as appropriate (i.e., received saline treatment). Mice were exposed to 2 cycles of CIE vapor. During acute withdrawal from CIE (4-6 hour) mice received CNO (3 mg/kg, I.P.) or saline in their home cage 30-min prior to the start of EPM testing (**Figure 6-7**).

**Figure 6.**
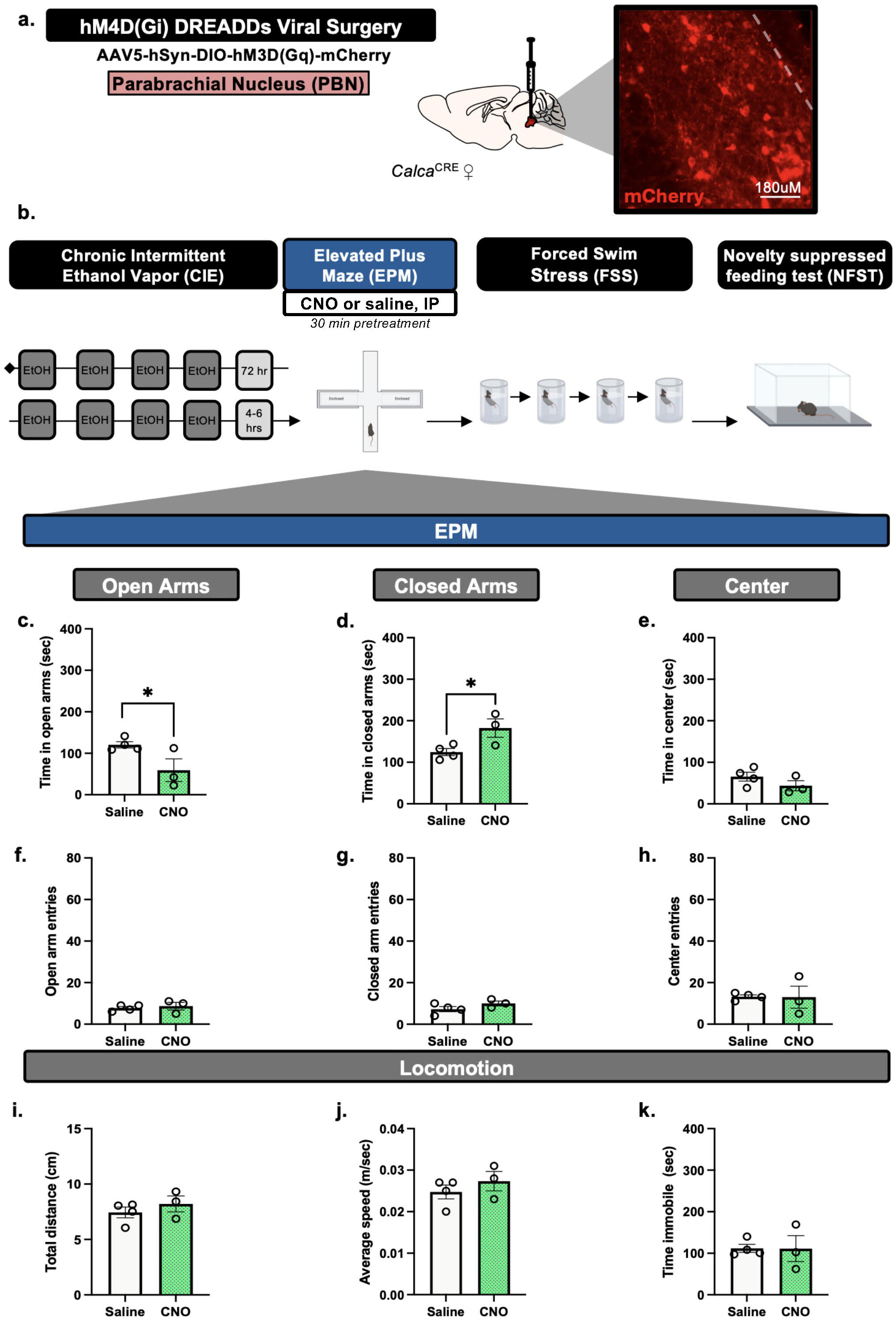
PBN(CGRP) inactivation in females during acute withdrawal from CIE alters time spent in the EPM arms. **a)** *Calca*^Cre^ female mice received bilateral viral infusions to express CRE-dependent hM4D(Gi) DREADDs in the PBN. Inset demonstrates representative image of hM4D(Gi) (mCherry) autofluorescence in the PBN. **b**) Mice received 2 cycles of CIE vapor exposure (one cycle = 4 days of 16 hrs of vapor, depicted in dark gray) with a 72 hr withdrawal between cycles. Next, during acute withdrawal (4-6 hrs post-final CIE exposure), mice were tested with EPM post-30 min pretreatment of CNO (3mg/kg, IP). **c**) CNO decreased time in open arms and **d**) increased time in closed arms compared to saline. **e**) CNO did not change time in center, **f**) open arm entries, **g**) closed arm entries, **h**) center entries, **i**) distance travelled, **j**) average speed or **k**) time immobile compared to saline. Values on graphs represent mean ± S.E.M., unpaired *t*-test (*p≦0.05).

**Figure 7.**
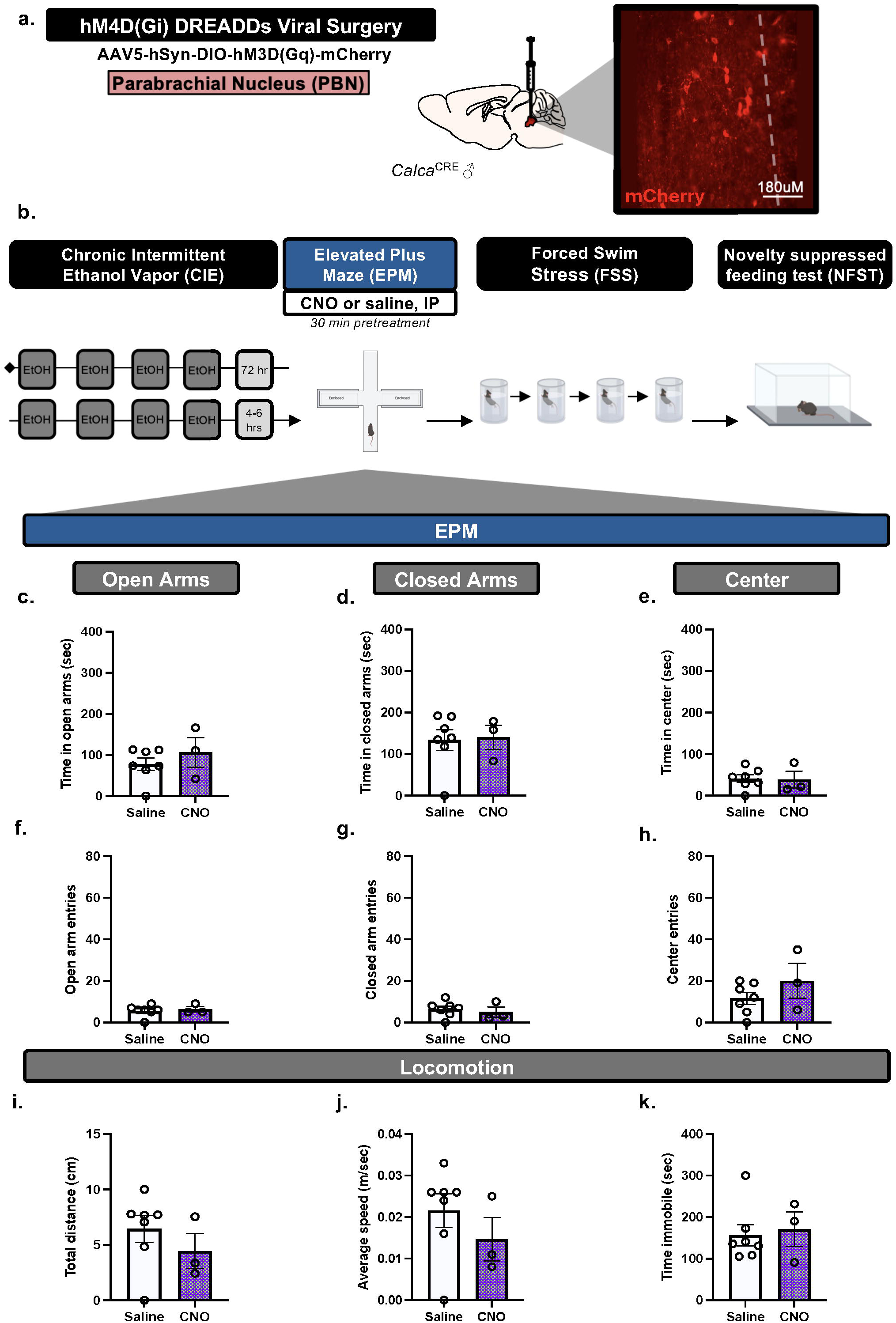
PBN(CGRP) inactivation in males during acute withdrawal from CIE does not change behavior in EPM. **a)** *Calca*^Cre^ male mice received bilateral viral infusions to express CRE-dependent hM4D(Gi) DREADDs in the PBN. Inset demonstrates representative image of hM4D(Gi) (mCherry) autofluorescence in the PBN. **b**) Mice received 2 cycles (one cycle = 4 days of 16 hrs on followed by 4 hrs of withdrawal) of CIE vapor exposure. Next, during acute withdrawal (4-6 hrs post-final CIE exposure), mice were tested with EPM post-30 min pretreatment of CNO (3mg/kg, IP). **c**) CNO did not change time in open arms, **d**) time in closed arms, **e**) time in center, **f**) open arm entries, **g**) closed arm entries, **h**) center entries, **i**) distance travelled, **j**) average speed or **k**) time immobile in EPM compared to saline. Values on graphs represent mean ± S.E.M., unpaired *t*-test (*p≦0.05).

#### 3.6.5. Experiment 5: Effects of PBN(CGRP) inactivation on NSFT following repeated FSS during prolonged CIE withdrawal

1 week into withdrawal from CIE, *Calca*^CRE^ female and male mice from Experiment 4 underwent FSS, previously described in (23). Mice were exposed as follows: 2 days of FSS, 2-day incubation period (i.e., no FSS), and 2 days of FSS. Mice received CNO (3 mg/kg, I.P.) or saline in their home cage 30-min prior to the each FSS (**Figure 8**). Treatment assignments remained consistent with Experiment 4. After the final FSS, all mice underwent the 3-day NSFT paradigm (**Figures 9-10**). No CNO or saline treatment was given prior to NSFT.

**Figure 8.**
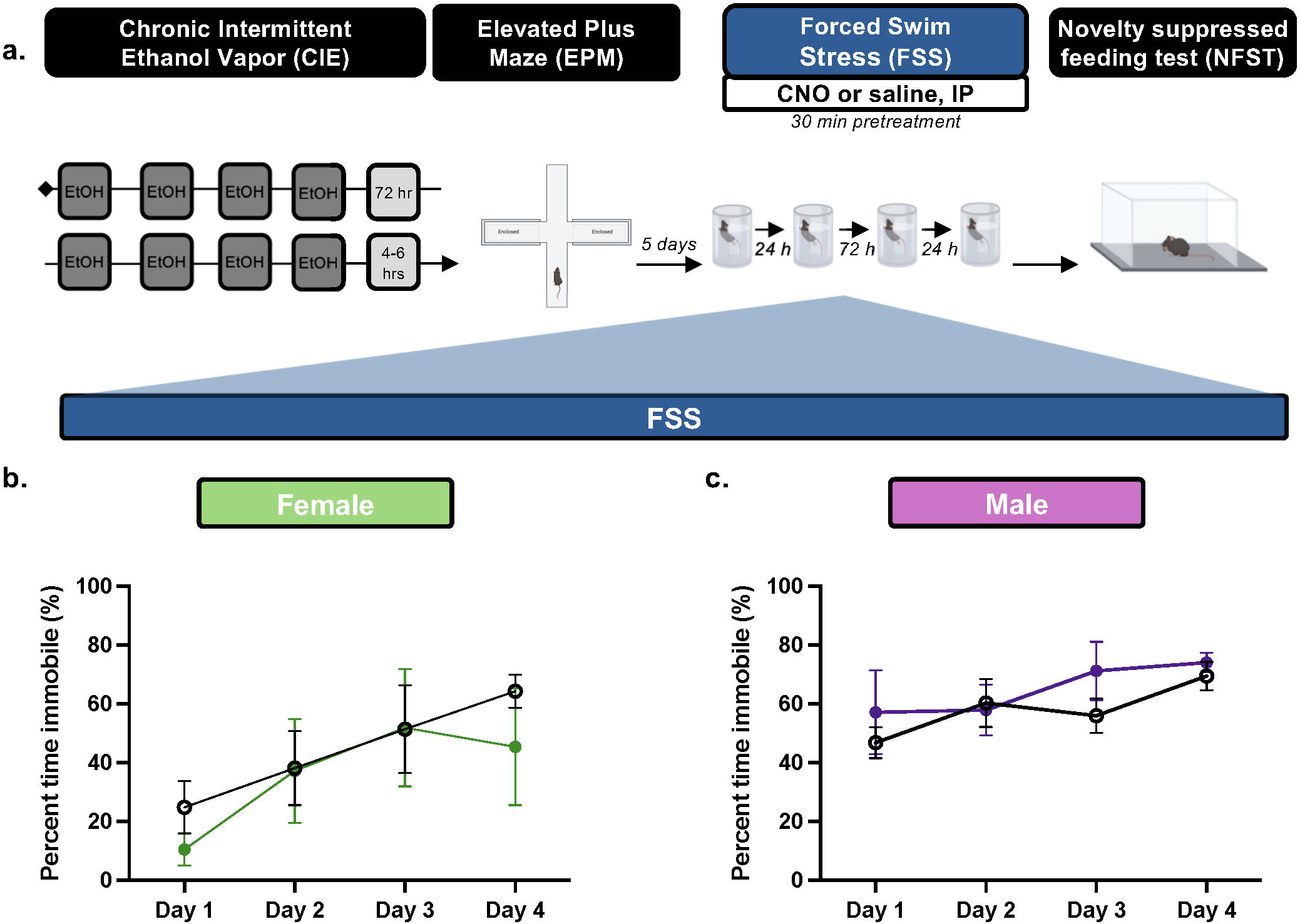
PBN(CGRP) inactivation during prolonged withdrawal from CIE does not change time immobile in FSS in females or males. **a)** 1 wk post CIE (i.e., 1 week after EPM), *Calca*^Cre^ male and female mice expressing CRE-dependent hM4D(Gi) DREADDs in the PBN received CNO (3 mg/kg, IP) or saline 30 min prior to a 6 min forced swim stress (FSS) for 4 days. **b**) In females, percent time immobile similarly increased across time with saline and CNO treatment, demonstrated by a significant main effect of day. **c**) In males, percent time immobile did not change across time with saline and CNO treatment. Values on graphs represent mean ± S.E.M., RM 2-way ANOVA (p≦0.05).

**Figure 9.**
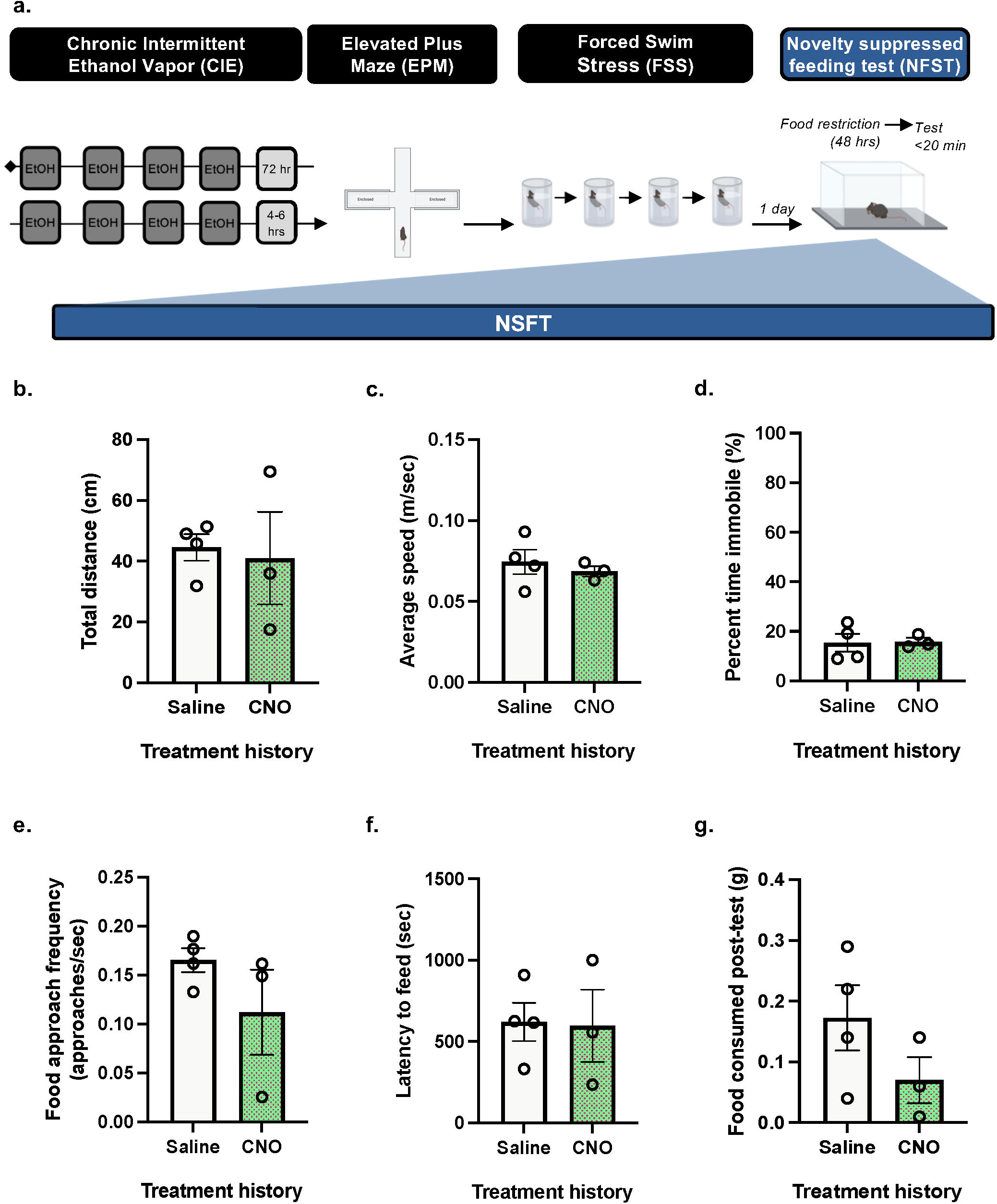
In prolonged withdrawal from CIE, PBN(CGRP) inactivation during FSS does not alter subsequent behavior in NSFT in females. **a)** 2 wks post CIE*, Calca*^Cre^ female mice with a history of CNO or saline treatment during FSS, were tested with NSFT. No chemogenetic manipulations occurred during NSFT. **b**) A history of CNO+FSS did not change distance travelled, **c**) average speed, **d**) percent time immobile, **e**) approach frequency, **f**) latency to feed, **g**) or food consumed post-test compared to a history of FSS+saline. Values on graphs represent mean ± S.E.M., unpaired *t*-test (*p≦0.05).

**Figure 10.**
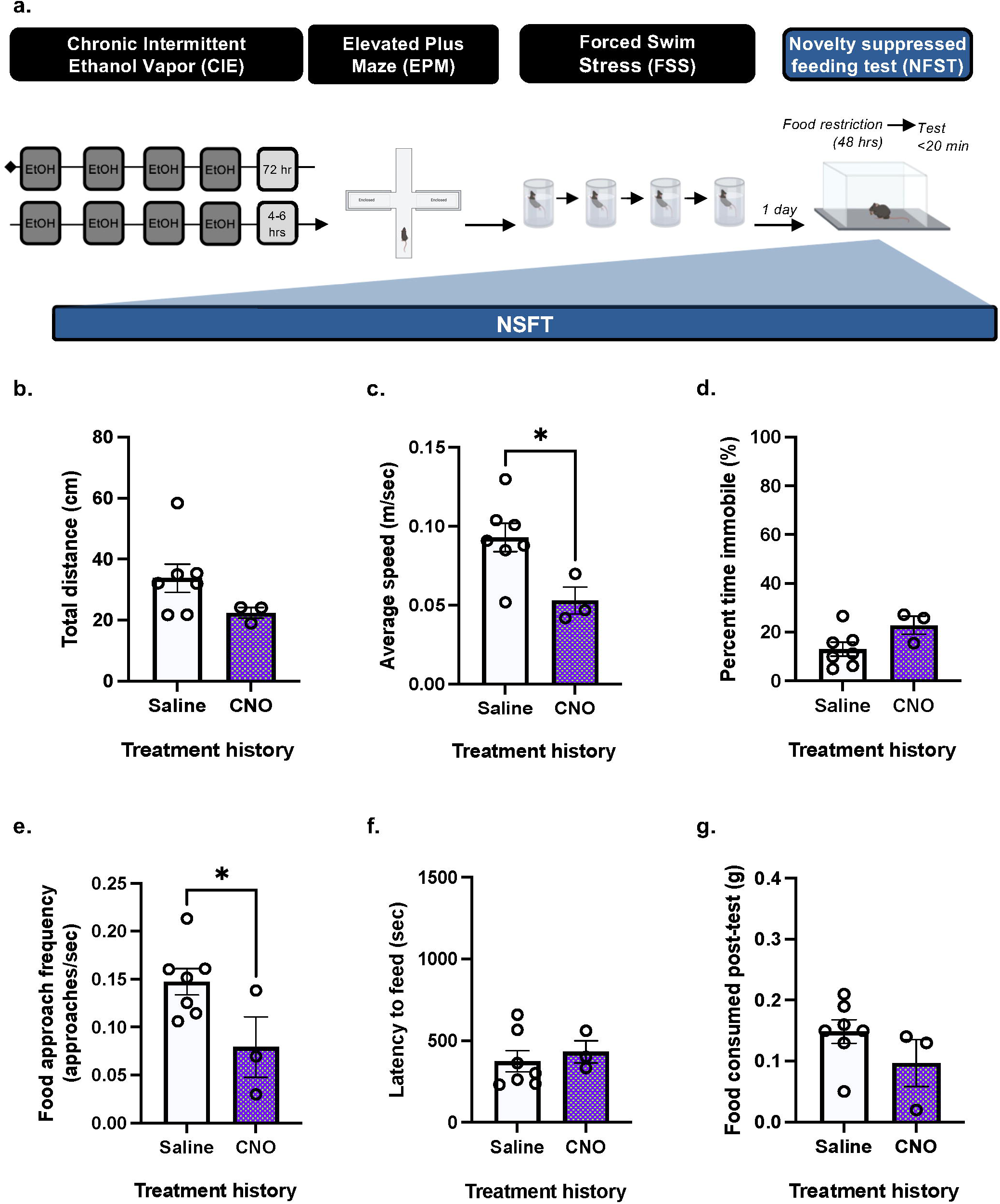
In prolonged withdrawal from CIE, PBN(CGRP) inactivation during FSS decreased subsequent approach frequency and speed in NSFT in males. **a)** 2 wks post CIE*, Calca*^Cre^ male mice with a history of CNO or saline treatment during FSS, were tested with NSFT. No chemogenetic manipulations occurred during NSFT. **b**) Distance travelled was similar in mice with a history of FSS+CNO relative to a history of FSS+saline. **c**) A history of FSS+CNO decreased average speed compared to a history of FSS+saline. **d**) Percent time immobile was similar in mice with a history of FSS+CNO relative to a history of FSS+saline. **e**) A history of FSS+CNO decreased approach frequency compared to a history of FSS+saline. **e**) Latency to feed in NSFT and **g**) food consumed post-test was similar in mice with a history of FSS+CNO relative to a history of FSS+saline. Values on graphs represent mean ± S.E.M., unpaired *t*-test (*p≦0.05).

### 3.7. Drugs

Sterile saline or 3 mg/kg Clozapine-N-oxide (CNO, MilliporeSigma) diluted in saline was delivered I.P. Pyrazole + Ethanol (1 mmol/kg + 1.6 g/kg Ethanol, IP) or Pyrazole (1 mmol/kg, IP) was diluted in sterile saline prior to each exposure.

### 3.8. Statistical analysis

Data is represented as mean ± S.E.M and all statistics were run using Prism 10 (GraphPad). Differences between groups were assessed using unpaired *t*-tests, paired *t*-tests, Repeated Measures (RM) Two-way analysis of variance (ANOVA), and Tukey’s post hoc tests, with significance set at p ≦ 0.05.

## 4. Results

### 4.1. Experiment 1: PBN(CGRP) activation does not alter behaviors in EPM in either females or males

To assess whether activating PBN(CGRP) neurons modulates behavior in EPM, *Calca*^CRE^ female and male mice received bilateral viral injections of CRE-dependent hM3D(Gq) DREADDs virus in the PBN (**Figure 1a**, **Figure 2a**). CNO (3 mg/kg, IP) or vehicle (saline) treatment was administered 30-min prior to EPM. In females, unpaired *t*-tests of total time spent in areas of the EPM demonstrate no significant effect of CNO relative to saline on time spent in open arms (t_7_=0.8008, p=0.4496; **Figure 1b**), closed arms (t_7_=1.035, p=0.3352; **Figure 1c**), or the center zone (t_7_=0.2324, p=0.8229; **Figure 1d**). Therefore, hM3D(Gq) activation by CNO did not affect time spent in the EPM areas. Unpaired *t*-tests of number of entries into the areas of the EPM demonstrate no significant effect of CNO relative to saline on open arm entries (t_7_=0.4637, p=0.6569; **Figure 1e**), a trend toward a significant effect on closed arm entries (t_7_=2.235, p=0.0605; **Figure 1f**), and no significant effect on center entries (t_7_=1.564, p=0.1618; **Figure 1g**). Therefore, hM3D(Gq) activation by CNO did not affect number of entries into the EPM areas. Unpaired *t-*tests of locomotor behaviors in the EPM demonstrate no significant effect of CNO relative to saline on total distance travelled (t_7_=1.268, p=0.2455; **Figure 1h**) or average speed (t_7_=1.687, p=0.1355; **Figure 1i**), and a trend toward a significant effect on total time immobile (t_7_=2.150, p=0.0686; **Figure 1j**). Therefore, hM3D(Gq) activation by CNO did not affect locomotor behaviors in the EPM. Overall, in females, activating excitatory hM3D(Gq) DREADDs expressed in the PBN via CNO did not change behaviors in EPM.

In males, unpaired *t*-tests of total time spent in areas of the EPM demonstrate no significant effect of CNO relative to saline on time spent in open arms (t_5_=0.3804, p=0.7193; **Figure 2b**), closed arms (t_5_=0.6773, p=0.5282; **Figure 2c**) or center zone (t_5_=0.5814, p=0.5862; **Figure 2d**). Therefore, hM3D(Gq) activation by CNO did not affect time spent in the EPM areas. Unpaired *t*-tests of number of entries into the areas of the EPM demonstrate no significant effect of CNO relative to saline on open arm entries (t_5_=1.320, p=0.2441; **Figure 2e**), closed arm entries (t_5_=0.07704, p=0.9416; **Figure 2f**), or center entries (t_5_=1.030, p=0.3504; **Figure 2g**). Therefore, hM3D(Gq) activation by CNO did not affect number of entries into the EPM areas. Unpaired *t-*tests of locomotor behaviors in the EPM demonstrate CNO relative to saline had no significant effect on total distance travelled (t_5_=0.4405, p=0.6780; **Figure 2h**), average speed (t_5_=0.4437, p=0.6758; **Figure 2i**), or total time immobile (t_5_=0.6721, p=0.5313; **Figure 2j**). Therefore, hM3D(Gq) activation by CNO did not affect locomotor behaviors in the EPM. Thus, in males, activating excitatory hM3D(Gq) DREADDs expressed in the PBN via CNO did not change behaviors in EPM.

### 4.2. Experiment 2: BNST^PBN^ activation does not alter behaviors in EPM in either females or <u>males</u>

To assess whether activating of BNST^PBN^ neurons modulates behavior in EPM, we used an anterograde transsynaptic viral transfer strategy (33–35) that would target PBN-innervated BNST cells. C57BL/6J female and male mice received bilateral injections of anterograde transsynaptic CRE virus in the PBN and CRE-dependent hM3D(Gq) DREADDs virus in the BNST (**Figure 3a**, **Figure 4a**). CNO (3 mg/kg, IP) or vehicle (saline) treatment was administered 30-min prior to EPM. In females, paired *t*-tests of total time spent in areas of the EPM demonstrate no significant effect of CNO relative to saline on time spent in open arms (t_4_=0.2459, p=0.8179; **Figure 3b**), closed arms (t_4_=0.05269, p=0.9605; **Figure 3c**), or the center zone (t_4_=1.392, p=0.2363; **Figure 3d**). Therefore, hM3D(Gq) activation by CNO did not affect time spent in the EPM areas. Paired *t*-tests of number of entries into the areas of the EPM demonstrate no significant effect of CNO relative to saline on open arm entries (t_4_=0.7845, p=0.4766; **Figure 3e**), closed arm entries (t_4_=1.554, p=0.1951; **Figure 3f**), or center entries (t_4_=1.411, p=0.2312; **Figure 3g**). Therefore, hM3D(Gq) activation by CNO did not affect number of entries into the EPM areas. Paired *t-*tests of locomotor behaviors in the EPM demonstrate a trend toward a significant effect of CNO relative to saline on total distance travelled (t_4_=2.659, p=0.0564; **Figure 3h**), and no significant effect on average speed (t_4_=2.539, p=0.0641; **Figure 3i**), or total time immobile (t_4_=2.030, p=0.1122; **Figure 3j**). Therefore, hM3D(Gq) activation by CNO did not affect locomotor behaviors in the EPM. Overall, in females, activating excitatory hM3D(Gq) DREADDs, expressed in BNST^PBN^ neurons, via CNO did not change behaviors in EPM.

In males, paired *t*-tests of total time spent in areas of the EPM demonstrate no significant effect of CNO relative to saline on time spent in open arms (t_6_=0.1034, p=0.9210; **Figure 4b**), closed arms (t_6_=0.3312, p=0.7518; **Figure 4c**) or center zone (t_6_=0.3960, p=0.7058; **Figure 4d**). Therefore, hM3D(Gq) activation by CNO did not affect time spent in the EPM areas. Paired *t*-tests of number of entries into the areas of the EPM demonstrate no significant effect of CNO relative to saline on open arm entries (t_6_=1.732, p=0.1340; **Figure 4e**), closed arm entries (t_6_=0.1709, p=0.8699; **Figure 4f**), or center entries (t_5_=1.6124, p=0.5628; **Figure 4g**). Therefore, hM3D(Gq) activation by CNO did not affect number of entries into the EPM areas. Paired *t-*tests of locomotor behaviors in the EPM demonstrate CNO relative to saline had no significant effect on total distance travelled (t_6_=0.7197, p=0.4988; **Figure 4h**), average speed (t_6_=0.6951, p=0.5130; **Figure 4i**), or total time immobile (t_6_=0.6850, p=0.5189; **Figure 4j**). Therefore, hM3D(Gq) activation by CNO did not affect locomotor behaviors in the EPM. Overall, in males, activating excitatory hM3D(Gq) DREADDs, expressed in BNST^PBN^ neurons, via CNO did not change behaviors in EPM.

### 4.3. Experiment 3: CGRP+ cell expression in the PBN and CGRP+ innervated cells in the BNST do not differ between sexes

To assess whether CGRP-containing cells in the PBN and CGRP-innervated cells in the BNST differed between the sexes, the PBN (**Figure 5a**) and BNST (**Figure 5c**) from CGRP-DTR^GFP^ female and male mice were analyzed for CGRP expression. Unpaired *t*-test of total colocalized CGRP (GFP) and DAPI cells in PBN slices demonstrated no significant difference in females relative to males (t_10_=0.9321, p=0.3732; **Figure 5b**). Similarly, unpaired *t*-test of total colocalized CGRP (GFP) innervated DAPI cells in PBN slices demonstrated no significant difference in females relative to males (t_10_=0.07935, p=0.9383; **Figure 5d**). Overall, the baseline quantity of CGRP-containing PBN cells or CGRP-innervated cells does not differ between females and males.

### 4.4. Experiment 4: PBN(CGRP) inactivation during acute alcohol withdrawal alters EPM behavior in females

To assess whether inactivating PBN(CGRP) neurons modulates EPM behavior during early withdrawal from CIE, *Calca*^CRE^ female and male mice received bilateral injections of CRE-dependent hM4D(Gi) DREADDs virus in the PBN (**Figure 6a**; **Figure 7a**). Mice underwent 2 cycles of CIE vapor exposure. During acute withdrawal (4-6 hrs post final CIE exposure), CNO (3 mg/kg, IP) or vehicle (saline) treatment was administered 30-min prior to EPM (**Figure 6b**; **Figure 7b**). In CIE-exposed females, unpaired *t*-test of total time spent in areas of the EPM demonstrate CNO relative to saline had a significant effect on time spent in open arms (t_5_=2.501, p=0.0544; **Figure 6c**), and a significant effect on time spent in closed arms (t_5_=2.736, p=0.0410; **Figure 6d**), with no significant effect on time spent in the center zone (t_5_=1.345, p=0.2365; **Figure 6e**). Therefore, hM4D(Gi) activation by CNO decreased time spend in the open arms and increased time spent in the closed arms compared to saline while not increasing time in the center. Unpaired *t*-tests of number of entries into the EPM areas demonstrate CNO compared to saline had no significant effect on open arm entries (t_5_=0.5125, p=0.6301; **Figure 6f**), closed arm entries (t_5_=1.557, p=0.1803; **Figure 6g**) or center entries (t_5_=0.05505, p=0.9582; **Figure 6h**). Therefore, hM4D(Gi) activation by CNO did not affect number of entries into EPM areas. Unpaired *t*-tests of locomotor behaviors in the EPM demonstrate CNO relative to saline had no significant effect on total distance travelled (t_5_=0.9222, p=0.3988; **Figure 6i**), average speed (t_5_=0.9351, p=0.3927; **Figure 6j**), or total time immobile (t_5_=0.01838, p=0.9860; **Figure 6k**). Therefore, hM4D(Gi) activation by CNO did not affect locomotor behaviors in the EPM. Overall, in females activating the inhibitory hM4D(Gi) DREADDs in the PBN, via CNO during early CIE-withdrawal, selectively decreased time spent in the open arms and increased time spent in the closed arms of the EPM.

In CIE-exposed males, unpaired *t*-tests of total time spent in areas of the EPM demonstrated CNO relative to saline did not significantly affect time spent in open arms (t_8_=0.8957, p=0.3966; **Figure 7c**), closed arms (t_8_=0.1478, p=0.8862; **Figure 7d**), or center zone (t_8_=0.1183, p=0.9087; **Figure 7e**). Therefore, hM4D(Gi) activation by CNO did not affect time spent in the EPM areas. Unpaired *t*-tests of number of entries into the EPM areas demonstrate CNO compared to saline had no significant effect on open arm entries (t_8_=0.4159, p=0.6884; **Figure 7f**), closed arm entries (t_8_=0.5307, p=0.6100; **Figure 7g**) or center entries (t_8_=1.260, p=0.2432; **Figure 7h**). Therefore, hM4D(Gi) activation by CNO did not affect number of entries into the EPM areas. Unpaired *t*-tests of locomotor behaviors in the EPM demonstrate CNO compared to saline had no significant effect on total distance travelled (t_8_=0.9334, p=0.3779; **Figure 7i**), average speed (t_8_=0.9671, p=0.3618; **Figure 7j**), or total time immobile (t_8_=0.3075, p=0.7663; **Figure 7k**). Therefore, hM4D(Gi) activation by CNO did not affect locomotor behaviors in the EPM. Thus, in males, activating the inhibitory hM4D(Gi) DREADDs in the PBN, via CNO during early CIE-withdrawal, did not alter behaviors in EPM.

### 4.5. Experiment 5: PBN(CGRP) inactivation in males during forced swim stress alters subsequent behavior in NSFT

Next, we assessed whether inactivating PBN(CGRP) neurons during repeated FSS exposure modulates subsequent NSFT behavior during prolonged withdrawal from CIE. On the 6^th^ day of withdrawal from CIE, *Calca*^CRE^ female and male mice from Experiment 4 were exposed to FSS for four days, with CNO (3 mg/kg, IP) or vehicle (saline) administered 30-min prior to testing (**Figure 8a**). Three days post-FSS, mice were tested on NSFT (**Figure 9a**, **10a**).

In females, two-way RM ANOVA of percent time immobile in FSS (**Figure 8b**) demonstrated a significant main effect of Time (F[3,15]=7.936, p=0.0035), no significant main effect of Treatment (F[1,5]=0.2677, p=0.6269), and no significant Time x Treatment interaction (F[3,15]=0.6536, p=0.5930). Therefore, in females, immobility was significantly increased across the 4 days of FSS exposure and did not differ between CNO and saline treatment. Conversely, in males, two-way ANOVA of percent time immobile in FSS (**Figure 8c**) demonstrated no significant main effect of Time (F[3,24]=2.980, p=0.0721), Treatment (F[1,8]=0.8061, p=0.3955) or Time x Treatment interaction (F[3,24]=0.6203, p=0.6087). Therefore, in males, time immobile did not change across sessions and did not differ between CNO and saline treatment. Overall, in females and males, activating the inhibitory hM4D(Gi) DREADDs did not alter repeated FSS behavior during prolonged CIE-withdrawal.

In FSS-exposed females, unpaired *t*-tests of locomotor behaviors in NSFT demonstrated no significant effect of a history of FSS + CNO relative to a history of FSS + saline on total distance travelled (t_5_=0.2592, p=0.8058; **Figure 9c**), average speed (t_5_=0.6204, p=0.5621; **Figure 9b**), or percent time immobile (t_5_=0.09971, p=0.9245; **Figure 9d**). Therefore, a history of hM4D(Gi) activation by CNO during FSS did not affect locomotor behaviors in the NSFT. Unpaired *t*-tests of food-associated behaviors in the NSFT demonstrated no significant effect of a history of FSS + CNO relative to a history of FSS + saline on approach frequency to food pellet zone (t_5_=1.361, p=0.2317; **Figure 9e**), latency to feed (t_5_=0.1009, p=0.9235; **Figure 9f**), or amount of food consumed post-testing (t_5_=1.443, p=0.2087; **Figure 9g**). Therefore, a history of hM4D(Gi) activation by CNO during FSS did not affect, number of approaches and final approach in the NSFT, or consummatory behaviors post-NSFT. Overall, in females, a history of activating the inhibitory hM4D(Gi) DREADDs expressed in the PBN, during the FSS sessions in prolonged CIE-withdrawal, did not alter behaviors in NSFT.

In FSS-exposed males, unpaired *t*-tests of locomotor behaviors in the NSFT demonstrated no significant effect of a history of FSS + CNO relative to a history of FSS + saline on total distance travelled (t_8_=1.535, p=0.1634; **Figure 10b**), a significant effect on average speed (t_8_=2.669, p=0.0284; **Figure 10c**), and no significant effect on percent time immobile (t_8_=1.974, p=0.0838; **Figure 10d**). Therefore, a history of activation of hM4D(Gi) by CNO during FSS did not globally affect locomotor behaviors (i.e., distance and immobility) but selectively decreased speed in the NSFT. Unpaired *t*-tests of food-associated behaviors in the NSFT demonstrated a significant effect of a history of FSS + CNO relative to a history of FSS + saline on approach frequency to food pellet zone (t_8_=2.365, p=0.0456; **Figure 10e**), and no significant effect on latency to feed (t_8_=0.5208, p=0.6166; **Figure 10f**), or food consumed post-testing (t_8_=1.358, p=0.2116; **Figure 10g**). Therefore, a history of hM4D(Gi) activation by CNO during FSS decreased approach frequency and did not affect final approach in the NSFT or consummatory behaviors post-NSFT. Overall, in males, a history of activating the inhibitory hM4D(Gi) DREADDs expressed in the PBN with CNO during the FSS sessions in prolonged CIE-withdrawal increased approach frequency to food pellet zone and decreased speed travelled in NSFT.

## 5. Discussion

Chemogenetically activating PBN(CGRP) or BNST^PBN^ neurons during EPM did not change behaviors under alcohol-naïve conditions for either sex. Experimentally-naïve females and males showed no sex differences in number of CGRP+ cells in the PBN and CGRP+ innervated cells in the BNST. Inhibiting PBN(CGRP) neurons in EPM during acute withdrawal from CIE increased anxiety-like behavior selectively in females. In the same cohort, inhibiting PBN(CGRP) neurons during repeated stress in prolonged withdrawal did not affect behavior but did subsequently increase anxiety-like behavior selectively in males. Thus, manipulating PBN(CGRP) neurons induced alcohol-, stress-, and sex-specific changes in anxiety-like measures in alcohol-withdrawal.

Previous work demonstrates activating PBN(CGRP) projections increases anxiety-like behaviors in both sexes. Specifically, optogenetically activating PBN(CGRP) neurons increased freezing behavior and decreased open arm time and preference in EPM in both sexes (19). Additionally, optogenetic stimulation of PBN(CGRP) projections also led to avoidance of the stimulation side of the real-time place-preference chamber (19). Similar anxiogenic effects were found with circuit-specific optogenetic activation of PBN to paraventricular thalamus projections in male mice, as anxiety-like behaviors increased in the open field test and increased avoidance of the photostimulation-paired chamber in real-time place aversion (36). Studies that pharmacologically manipulate CGRP also demonstrate an anxiogenic role, as intra-BNST infusions of CGRP agonist enhanced anxiety-like measures in EPM (13, 14). Given the PBN is the dominant extrinsic source of CGRP in the BNST (3, 4, 16–18), these studies indirectly implicate a role for the PBN(CGRP) ➜ BNST circuit in modulating anxiety. However, our findings demonstrate chemogenetic activation of PBN(CGRP) or BNST^PBN^ neurons does not change anxiety-like behaviors during EPM in either sex. The previous findings used optogenetic and pharmacological manipulations, which offer instant control of neuronal activity (37) and direct manipulation of an endogenous receptor, respectively. Given our use of DREADDs induces gradual neuromodulation through heighten excitability (37, 38), the optogenetic and pharmacological manipulations during the passive and active avoidance assays may elicit significant anxiety-like responses due to their robust ability to engage neuronal control. Relative to other studies that used chemogenetic manipulations, our findings suggest that that the behavioral assay used to investigate the anxiogenic role of PBN(CGRP) neurons may also result in different effects. When measuring anxiety-like behavior with NSFT, an approach-avoidance conflict task, in females and males, chemogenetically activating PBN(CGRP) neurons increases latency to feed and frequency of non-consummatory approach bouts (21). Circuit-specific chemogenetic BNST^PBN^ neuron activation also increases latency to feed albeit selectively in females (21). Given the difference in behavioral assays, (i.e. EPM compared to NSFT), chemogenetically activating PBN and BNST^PBN^ neurons may be more sensitive to anxiety-like behavior in approach-avoidance contexts. Altogether, our studies suggest that in naïve conditions increasing PBN(CGRP) neuron excitability is not sufficient to induce anxiety-like behavior in a passive avoidance context.

While no anxiety-like behaviors in EPM arose following chemogenetic PBN(CGRP) manipulation in alcohol-naïve mice, we demonstrate chemogenetically inhibiting PBN(CGRP) neurons during EPM in acute withdrawal from CIE increased anxiety-like behavior selectively in females. Thereby, the data suggest that decreasing PBN(CGRP) neuron excitability is sufficient to modulate anxiety-like behavior following alcohol exposure in a sex-specific manner. A history of adolescent alcohol vapor exposure shows a similar effect in adult females, as chemogenetic BNST activation increased latency to feed in the novelty-induced hypophagia test that was associated with increased cFOS activity in PBN(CGRP) neurons (28). Given the BNST sends projections to the PBN (3, 16), the findings suggest indirect activation of the PBN(CGRP) neurons via the BNST is associated with a heightened anxiety in prolonged alcohol-abstinence. Thereby, after alcohol exposure, females may show increased sensitivity to PBN(CGRP) manipulations (excitatory or inhibitory) that result in anxiety-like behavior, an effect not seen in alcohol-naïve conditions. The lack of effect in males during acute withdrawal is in line with previous work that demonstrates CGRP expression is upregulated in select brain regions only in prolonged withdrawal from CIE and not during exposure (27). Similarly, our later findings in prolonged withdrawal from CIE reveal that PBN(CGRP) inhibition during stress subsequently increases anxiety-like behavior. Thereby, alcohol-associated changes in the PBN(CGRP) system are present, albeit in select alcohol-contexts. Indeed, previous studies exclusively conducted in males demonstrate that alcohol-associated changes in the PBN(CGRP) system are present. In the BNST, CGRP terminal-expression increases in response to voluntary drinking in alcohol-preferring male rats (26), thereby suggesting heightened incoming PBN(CGRP) activity. Moreover, dysregulations in the PBN(CGRP) system in males may also be a genetic predisposition for alcohol-preference. CGRP receptor binding sites in alcohol-preferring and high alcohol-drinking male rats are significantly lower in the central amygdala (CeA; (25)), which predominately receives CGRP from PBN(CGRP) projections (3, 17). Future studies will need to investigate if the alcohol-related changes in CGRP expression in males are also present in females. Thus, in males the brain region-specific changes in CGRP expression occur at baseline in alcohol-preferring rats, with voluntary intake, and selectively in prolonged withdrawal from CIE. While PBN(CGRP)-induced changes in anxiogenic behavior occur in acute withdrawal in females and in prolonged withdrawal in males.

In addition to modulating anxiety, previous work shows that PBN(CGRP) neurons also modulate freezing and immobility in threat contexts (4). For example, ablating PBN(CGRP) neurons reduces freezing in response to foot shock (20). Similarly, optogenetically activating PBN(CGRP) neurons without external stimuli induces freezing and immobility behavior (19, 20, 22). Moreover, a history of FSS, an assay that also induces immobility behavior, is associated with dysregulated *ex vivo* PBN(CGRP) ➜ BNST activity (23). Therefore, we investigated if inhibiting PBN(CGRP) neurons would modulate immobility behavior in the FSS paradigm during prolonged withdrawal from CIE. We found chemogenetic inhibition of PBN(CGRP) neurons did not impact FSS immobility in withdrawal. Moreover, repeated PBN(CGRP) inhibition did not change immobility, as females continued to display increased immobility across days with no change in males. When taken with previous work that demonstrates a role for PBN(CGRP) in foot shock-induced freezing or immobility in the absence of a stimuli, our data using forced swim indicate manipulating PBN(CGRP) does not globally affect immobility responses in both sexes. Therefore, the role of PBN(CGRP) neurons in immobility may be context dependent such that it is limited to painful stimuli or threatening open field contexts.

Interestingly, despite PBN(CGRP) inhibition not affecting repeated FSS behavior during prolonged withdrawal in both sexes, the manipulations had subsequent effects on anxiety-like behavior selectively in males. Specifically, a history of chemogenetically inhibiting PBN(CGRP) neurons during FSS decreased approach frequency in NSFT, suggesting increased avoidance of the center, an anxiety-like phenotype. Speed was also decreased albeit there was no effect on distance travelled or immobility, suggesting that motor function was intact. Therefore, the behavioral changes induced by inhibition of PBN(CGRP) neurons in this approach-avoidance conflict assay implies a change in threat assessment. The role of PBN(CGRP) in avoidance has been shown with PBN(CGRP) manipulations in real time, as optogenetically activating PBN(CGRP) neurons during an actively threatening context (i.e. chasing robot) enhanced avoidance behaviors during conditioning and recall in both sexes (22). To our knowledge ours is the first study to demonstrate that manipulating PBN(CGRP) neurons during stress altered a subsequent anxiety-like behavior. The role of PBN(CGRP) neurons in modulating subsequent behavior is in line with its role in fear learning. For example, optogenetically activating PBN(CGRP) neurons induced immobility in the absence of external stimuli, and pairing optogenetic PBN(CGRP) activation with a cue or context resulted in conditioned freezing behavior (19, 20). Additionally, optogenetic photoinhibition or ablation of PBN(CGRP) neurons decreased freezing during a foot shock and subsequent cue or context conditioned freezing behavior (19, 20). While we did not test associative learning, manipulating PBN(CGRP) neurons during stress similarly affected a later outcome, with an anxiety-like context rather than fear-related. Thus, manipulating PBN(CGRP) neurons can generate later behavioral changes. Our previous study suggests that the subsequent effects on anxiety-like behavior may be in part due to neuroadaptations within the PBN(CGRP) ➜ BNST circuit (23). Using the same paradigm, FSS followed by NSFT, we demonstrated that in alcohol-naïve conditions repeated stress induces anxiety-like behavior that associated with dysregulated *ex vivo* PBN(CGRP) ➜ BNST activity in males (23). Our findings that inhibiting PBN(CGRP) neurons during stress increases anxiety-like behavior in withdrawal further implicates that stress– and anxiety-contexts recruit PBN(CGRP) neurons, which may be protective against the later behavioral effects of stress. A previous study supports the protective role of CGRP in stress, as intracerebroventricular CGRP administration prior to chronic stress exposure blocked stress-induced depressive-like behavior (8). Altogether, our data imply that males are sensitive to PBN(CGRP) manipulations during stress in prolonged withdrawal, and the recruitment of PBN(CGRP) neurons during stress exposure may be protective in a later approach-avoidance context.

Our findings indicating that PBN(CGRP) manipulations only had an effect in females during a passive avoidance context in acute withdrawal, and in males post-stress in an approach-avoidance context in prolonged withdrawal, suggest that PBN(CGRP) are differentially affected in both sexes. Interestingly, our findings in alcohol-naïve conditions show that there are no sex-specific differences in CGRP expression within the PBN neurons or fibers in the BNST. While not tested directly in our study, our behavioral findings suggest that alcohol exposure gives rise to sex-specific differences in PBN(CGRP) neuron expression and/or activity. Previous work supports this theory, as CGRP signaling show sex-specific changes in disease states. For example, women with knee osteoarthritis show higher CGRP mRNA expression in knee synovial compared to men with osteoarthritis (39). Others have shown in the acute phase of a neuropathic pain model, CGRP mRNA expression in the CeA was significantly upregulated selectively in male rats. However, in the chronic phase, CGRP and CGRP receptor component mRNA expression was significantly upregulated in females, whereas males did not show a significant increase (40). These data highlight that CGRP signaling in disease states are sex-specific and may change over time. Taken together, our data indicate alcohol-exposed females may be sensitive to CGRP manipulations during acute withdrawal. In contrast, alcohol-exposed males show sensitivity to CGRP manipulations after stress in prolonged withdrawal.

### 5.1. Conclusion

CGRP-target therapeutics are being considered for alcohol-induced headaches due to their efficacy in treating migraines (6, 7). Given that negative affect and stress can have detrimental effects during alcohol withdrawal in patients with AUD, the use of CGRP treatments in this context requires more research. Considering our findings, inhibiting CGRP-associated signaling may be disadvantageous for females in acute withdrawal, as it may be anxiogenic. While in males, chronically inhibiting CGRP-associated signaling during stress exposure may be anxiogenic in prolonged withdrawal. However, our data are specific to inhibition of PBN(CGRP) neurons, thus global CGRP inhibition may suppress or recruit signaling in such a way that negative affect is not worsened in AUD. These findings provide caution for the use of CGRP inhibition as a pharmaceutical treatment for AUD, while lending credence to the differential effects treatments can have on both sexes.

## Data Availability Statement

The data that support the findings of this study are available from the corresponding author upon reasonable request.

## Ethics Statement

The animal study was approved by Institutional Animal Care and Use Committee. The study was conducted in accordance with the local legislation and institutional requirements.

## Author Contributions

CVD: Investigation, Visualization, Writing-original draft, Writing-review & editing. JBT: Investigation, Writing-review & editing. AAJ: Conceptualization, Funding acquisition, Investigation, Methodology, Project administration, Supervision, Writing-original draft, Writing-review & editing.

## Acknowledgements

We thank Danny G. Winder (University of Massachusetts) for his support as a postdoctoral fellowship advisor and for providing the *Calca*^CRE^ and CGRP-DTR^GFP^ transgenic mouse lines. We thank Jill R. Turner (University of Kentucky) for access to behavioral and imaging resources. Graphics provided by BioRender (BioRender, Toronto, ON, Canada).

